# Feature engineering coupled machine learning algorithms for epileptic seizure forecasting from intracranial EEGs

**DOI:** 10.1101/131482

**Authors:** Rishav Kumar, Rishi Raj Singh Jhelumi, Achintye Madhav Singh, Prasoon Kumar

## Abstract

Epilepsy is one of the major neurological disorders affecting nearly 1 percentage of the global population. The major blunt is born by under developed and developing countries due to expensive treatment of epileptic conditions. Further, the lack of proper forecasting methods for an occurrence of epileptic seizures in epileptic-drug resistant patients or patients not amenable for surgery affects their psychological behaviour and restricts their daily activities. The forecasting is usually performed by human experts that leave a wide gap for human-bias and human error. Therefore, in the current work, we have evaluated the efficiency of several machine learning algorithms to automatically identify the preictal patterns corresponding to epileptic seizures from intracranial EEG signals. The robustness of the machine learning algorithms were tested after the data set was pre-processed using carefully chosen feature engineering strategies viz. denoised Fourier transforms as well as cross-correlation across electrodes in time and frequency domain. Extensive experimentations were carried out to determine the best combination of feature engineering techniques and machine learning algorithms. The best combination of feature engineering techniques and machine learning algorithm resulted in 0.7685 AUC (Area under the Receiver Operating Characteristic curve) on the random test samples. The suggested approach was fairly good at prediction of epilepsy in random samples and therefore, it can be used in epileptic seizure forecasting in patients where medication/surgery is ineffective. Eventually, our strategy reveals a robust method for brain disorders forecasting from EEGs.

## 1. Introduction

One of the most common neurological disorders, epilepsy, afflicts nearly 50 million people worldwide. It is a disorder which is marked by a sudden abnormal activity of a brain, characterised by seizures. Although, even after several type of drug-based medication and resective epileptic surgery, about 20-30% of people still suffers from epilepsy. A major reason behind reoccurrence of epilepsy is the development of drug-resistant epilepsy. Further, resective surgery in which a portion of brain is surgically removed in drug-resistant patients cannot be performed in every patient. The constant fear of reoccurrence of epilepsy in patients limits their participation in daily activities and has significant psychological impact on these patients[1]. So, there is a need to develop better seizure treatment management strategies where patients can be cautioned before-hand about the upcoming attack by seizures. This will enable them to get prepared for such risky situation and take effective medication/treatment for their well-being.

The most common diagnostic tool for an identification of an epileptic seizure is electroencephalography (EEG). The EEG signal signature is captured by tiny electrodes placed at several locations on the scalp of the patient’s brain. These electrodes record the voltage fluctuation due to the ionic current traversing along the length of neurons in the brain. Thus, any abnormal voltage recording during EEG measurement can be a potential signature of seizures in comparison to the normal signatures of the brain activity. EEG can be utilized to differentiate between epileptic and non-epileptic seizures through and locate the pattern corresponding to ‘interictal epileptiform discharges’ prevalent in epileptic patients with the help of experts. EEG can also be used to differentiate between different types of epileptic syndrome. However, developing a tool based on EEG analysis for early warning of seizures requires identification of pattern in EEG signals that precedes the actual seizure signals from epileptic patients. The reports on the existence of consistent signature of local field potential (LFP) before any seizure attack in patients can be used as potential tool for identifying the brain state and predict upcoming seizures [2]. Numerous clinical studies on epileptic patients have statistically confirmed the presence of seizure prone states, hours or days before the actual seizure attack. Moreover, the changes in cortical excitability, cerebral blood flow and its oxygenation are also prominent markers that have been measured preceding seizures.

The algorithm developed for an identification of the seizure signatures from EEGs have been developed in the past; however, they suffer from majorly from statistical rigor required for an effective prediction[5][7] Nevertheless, some attempts have recently been able to present the desirable statistical rigor for effective prediction of epilepsy[6] The major difficulty in the development of effective seizure predicting algorithm has been the scarcity of long recordings of EEG having ample signals of seizures, better featuring engineering for the extraction of patterns amidst noisy signal and implementation of better prediction algorithm. Machine learning has come along a long way for the pattern identification purpose and several algorithms have been implemented for seizure prediction in EEG patterns to identify brain disorders.

In the current work, we have studied best feature extraction strategies from the intracranial EEG dataset so that only relevant information is supplied to prediction computational algorithms. Feature engineering consisted of denoised fourier transform as well as multiple cross correlation between data from different devices. We passed the filtered feature set to several standard machine learning techniques. We experimented with different parameters of the computational classifiers and their combination with feature set to determine the best combination to yield efficient forecasting results.

## 2. Materials and Methods

### 2.1. Materials

The data set of the electroencephalogram (EEG) of 5 dogs were obtained from International Epilepsy Electrophysiology portal (www.ieee.org) funded by National Institute of Health (NIH). The algorithms for processing of data were written and tested through signal processing and Machine learning toolbox of MATLAB 7.4 on Dell Inspiron Laptop having Intel i5 four core processor with 4GB RAM. The all the codes were run on single core with no multithreading.

### 2.2. Methods

#### 2.2.1. Collection and sorting of EEG data

Sufficiently large EEG data sets were pulled out of International Epilepsy Electrophysiology portal to train our computational algorithm to detect and learn the changes in EEG signals during epileptic seizures. Care was taken to ensure that the EEG data has the preictal stage as well as interictal stage for the machine learning algorithms to learn the difference between either of the two. Intracranial EEG data procured from International Epilepsy Electrophysiology online portal was obtained by taking recordings for 5 dogs, which were afflicted by epilepsy using an ambulatory monitoring system [2, 4]. The EEG was collected from 16 electrodes from different parts of brain and each of them was sampled at 400 Hz. The subsequent voltage values were referenced using a group average method. These recordings over a good time span were able to contain close to 100 seizures as well. Apart from the canine patients’ EEG, EEG from 2 human epileptic patients was also obtained. However, for the latter case, the number of electrodes varied (up to 24) and the sampling were performed at 5000 Hz. Voltage referencing was done using an electrode outside the brain. The whole recordings amounted to 2500 hours in the final dataset obtained. The whole process is illustrated in Figure 1[4] which was obtained from kaggle.com.

**Figure 1:**
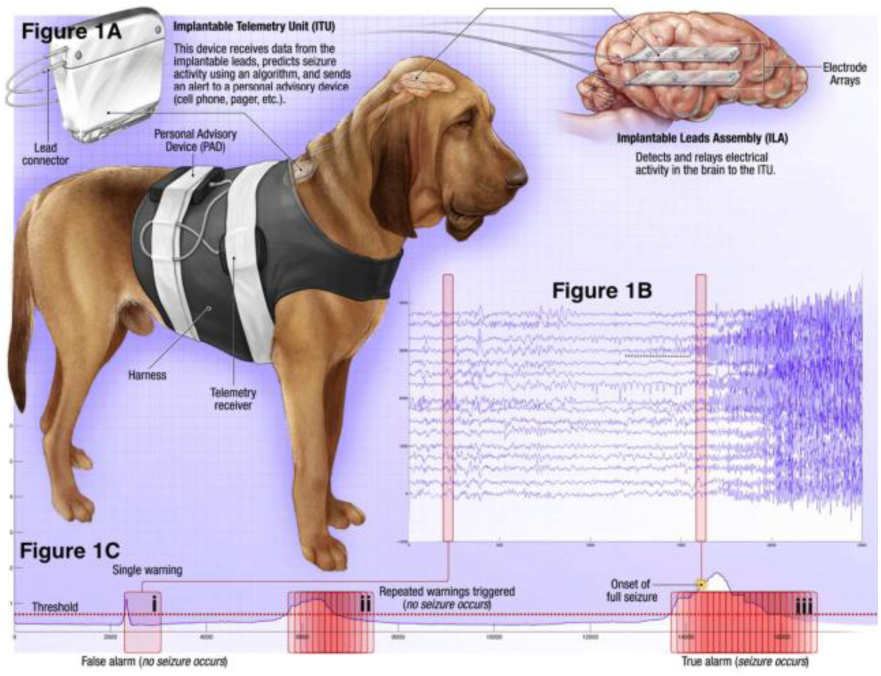
A) shows the way the intracranial EEG was recorded from canine subjects B) shows the EEG waves containing preictal phase and octal phase C) shows the sample working of any algorithm to work for correct prediction of seizures [4].

Based on the above EEG recordings, strategies were developed to differentiate whether a ten minute long EEG recording clip belongs to preictal phase or interictal phase. The following restrictions were placed to the definitions of preictal phase and to the interictal training data clip to avoid seizure clustering detection [8]

□ A data clip labelled as preictal phase will always be in the time range from one hour prior to the seizure to the onset of seizure.
□ A data clip labelled as interictal phase will always be separated by one week with any seizure in case of canine dataset and four hours in case of human dataset.

The final dataset that was obtained from above procedure is described as follow. With respect to training data, for every subject from the available five canine and two human subjects, *x* instances of 1 hour inter-ictal recordings and *y* instances of 1 hour pre-ictal recordings were obtained where *x* > *y*. Every such instances were divided into six ten minute data clips which were labelled by the ordered sequence number. The unlabelled testing data consisted of 900 ten-minute data clip to be classified by the computational algorithms.

A sample ten minute data clip consisted of two dimensional matrix of EEG sample values arranged row x column as electrode x time. With the modality of the data being time-series, there were close to 239766 voltage values per electrode in every ten minute data clip making the dimensionality of the data very high. The whole data set was 120 GB in size.

#### 2.2.2. Feature engineering strategies

As the dimensionality of data was huge (∼240,000), there was a need to extract relevant information or features on which we could run the machine learning techniques. This required the domain knowledge of large scale signal analysis and processing.

##### Method A

First, the recorded voltages were referenced to the group voltage to remove device errors. Second, Fourier transform of the data was computed to determine the major frequencies contributing to the relevant information. Fig 2(a) shows the time domain plot of interictal wave and (b) shows the frequency domain plot of interictal wave. Fig 3(a) shows the time domain plot of preictal wave and (b) shows the frequency domain plot of preictal wave. Third, we applied a low pass filter to remove the higher frequency component which added no significant information to the wave. The cut-off chosen was using hit and trial method which yielded the best results. Fourth, Power spectral density was calculated for this Fourier transform.[3] Fifth, to reduce the dimensionality, mean was computed over sequential rectangular windows. To arrive at the optimum length of rectangular window, we calculated standard deviation over different lengths of window and arrived at a value which yielded minimum standard deviation. On this feature data we applied different machine learning classifiers to classify.

**Figure 2.**
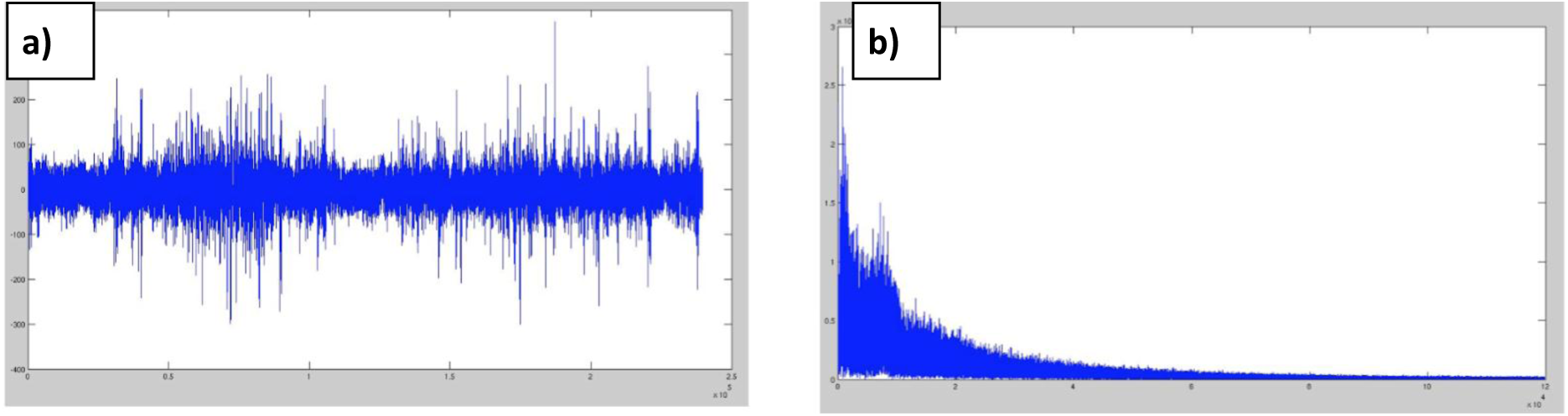
a) Time series plot of interictal wave and b) Fourier transform of interictal wave.

**Figure 3.**
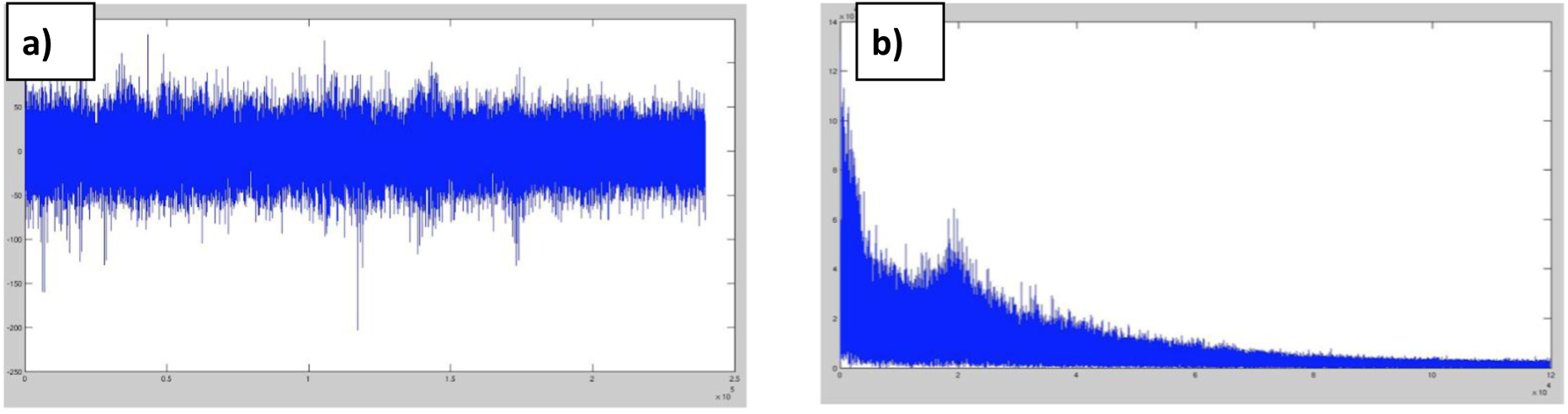
a) Time series plot of preictal wave and b) Fourier transform of preictal wave.

##### Method B

To get a factor of dependence between the electrodes signifying the ways in which different parts of the brain are interconnected, we followed a different approach. First, cross-correlation of the waves across different electrodes in time space was computed yielding n x n matrix where n is the number of electrodes.

Second, cross-correlation of the waves across different electrodes in frequency space was computed yielding n x n matrix where n is the number of electrodes. Third, eigenvector of the correlation was computed. Fourth, the entire above separate feature sets were concatenated to yield the final feature set on which we applied different machine learning classifiers to classify.

#### 2.2.3. Machine learning algorithms

After the feature set extraction standard machine learning algorithms were applied with different set of parameters and their AUC scores were compared to choose the best technique.

Extensive experimentation with different probabilistic classifiers was done. Chiefly those classifiers were:

□ SVM (kernels : linear, higher degree polynomial, radial)
□ Neural networks (different number of nodes and hidden layers)
□ Random Forests (number of trees : 1000, 3000, 10000)
□ Linear regression

After several experiments, we did a comparative study between different techniques with different feature sets which we will discuss in the results section. Linear regression performed the worst among all of them and we did not include in the comparative study.

## 3. Results and discussion

The diagnosis of epilepsy or the ability to forecast epileptic seizures is typically done by observing the different changes the brain undergoes during the onset of seizure or during the seizure itself. The electroencephalogram (EEG) of the brain is one of the ways to detect such changes. The EEG usually shows waves having a lot spikes with rapid changes during the onset of seizures which continues in the seizure as well. This change in pattern of the brain waves can be picked up by computational algorithms to efficiently forecast an impending seizure with some probability.

The different probabilistic classifiers were run on the testing data clips and accordingly classified as pre-ictal and interictal stage. We tried different probabilistic classifiers with different feature set to determine the best classifier-feature set combination to forecast the seizure

**Table 1.** shows the results with 8 different machine learning techniques. The metric we used to compare different techniques was AUC (Area under the ROC curve) score.

| Sl. No. | Classifier | Feature Set | AUC Score |
| --- | --- | --- | --- |
| 1. | NN : 1 hidden layer (100) | FFT + L.P.F + Mean(6%, 48) | 0.71084 |
| 2. | NN : 5 hidden layer (10 each) | FFT + L.P.F + Mean(6%, 48) | 0.70786 |
| 3. | NN : 1 hidden layer (100) | FFT + L.P.F + Mean(3%, 24) | 0.68232 |
| 4. | NN : 1 hidden layer (100) | Time Correlation + FFT Correlation + eigenvector | 0.62381 |
| 5. | SVM (Degree 2 kernel) | FFT + L.P.F + Mean(6%, 48) | 0.76585 |
| 6. | SVM (Radial Bias Kernel) | FFT + L.P.F + Mean(3%, 24) | 0.54953 |
| 7. | SVM (Linear Kernel) | FFT + L.P.F + Mean(6%, 48) | 0.69909 |
| 8. | Random Forests (10,000) | Time Correlation + FFT Correlation + eigenvector | 0.60880 |

The first 4 classifiers are neural networks with variable hidden layers. The number in brackets represents the number of nodes in the hidden layers. The next three classifiers are Support Vector machines with different kernels or transforming functions. The degree-2 polynomial kernel is used in the first SVM. The last classifier is Random Forests where the number in brackets is the number of trees.

The feature set tried were FFT, denoising Low filter and Mean computation over a rectangular window. The numbers in the bracket are the low pass filter cutoff and the size of the rectangular window. Other feature set was time correlation and eigenvector. Based on the results, it was evident that SVM with a degree two polynomial kernel over a feature set of denoised fourier transformed EEG data yielded the best results. We achieved a AUC score of 0.76585 which roughly translates to strongly fair to weakly good prediction. Another interesting point of note was that the random forests consistently performed poor despite variable number of trees, the investigation of which is out of scope of this paper.

## 4. Conclusion

With this comparative study, we not only highlighted the success of machine learning techniques to forecast immediate epileptic seizures, but also with extensive experimentation found the best combination of learning techniques and feature set to yield the best prediction. Our method shows that machine learning is the way forward in epileptic seizure forecasting and should be further investigated into. Further research and development should be done in this direction so that we see a working prototype in future helping epileptic patients living a normal life.

## Acknowledgement

The authors wish to acknowledge following entities for their support: Kaggle.com-for highlighting the problem and providing a platform to test out the algorithms and Ieee.org-for being the sponsor for the above platform’s competition and providing the EEG dataset on which the algorithms were tested.

